# Pins & Needles: Towards Limb Disownership in Augmented Reality

**DOI:** 10.1101/349795

**Authors:** Oliver A Kannape, Ethan JT Smith, Peter Moseley, Mark P Roy, Bigna Lenggenhager

## Abstract

The seemingly stable construct of our bodily self depends on the continued, successful integration of multisensory feedback about our body, rather than its purely physical composition. Accordingly, pathological disruption of such neural processing is linked to striking alterations of the bodily self, ranging from limb misidentification to disownership, and even the desire to amputate a healthy limb. While previous embodiment research has relied on experimental setups using supernumerary limbs in variants of the Rubber Hand Illusion, we here used Augmented Reality to directly manipulate the feeling of ownership for one’s own, biological limb. Using a Head-Mounted Display, participants received visual feedback about their own arm, from an embodied first-person perspective. In a series of three studies, in independent cohorts, we altered embodiment by providing visuotactile feedback that could be synchronous (control condition) or asynchronous (400ms delay, Real Hand Illusion). During the illusion, participants reported a significant decrease in ownership of their own limb, along with a lowered sense of agency. Supporting the right-parietal body network, we found an increased illusion strength for the left upper limb as well as a modulation of the feeling of ownership during anodal transcranial direct current stimulation. Extending previous research, these findings demonstrate that a controlled, visuotactile conflict about one’s own limb can be used to directly and systematically modulate ownership – without a proxy. This not only corroborates the malleability of body representation but questions its permanence. These findings warrant further exploration of combined VR and neuromodulation therapies for disorders of the bodily self.

## 1. INTRODUCTION

The foundations of our “selves”, and our understanding of who we are, are laid by the continuous and successful integration of multisensory information about our body. This interdependence between “mind” and body has occupied the thoughts of scholars time and again: from Descartes’ notion that the self cannot exist without the senses (Descartes & Cottingham, 2013), to Husserl positing that there is no possibility of distancing the self from the body or the body from the self (Husserl, 2002), to William James’ oft-cited claim that the body is “always there” (James, 1890). There is an overwhelming sense that the direct neural representation of our bodies, in harmony with our actual body composition, forms the basis of an infallible bodily self. However, clinical examples challenge this notion and have suggested that the body indeed can be experienced as lost, not under control, or not belonging (Brugger & Lenggenhager, 2014). This latter feeling of ownership is argued to be a key aspect of our sense of a bodily self (Blanke, 2012). Yet, while psychological and neuroscientific research has extensively investigated the fundaments of the feeling of ownership for a foreign body part, the clinically presented loss of ownership has largely been neglected.

Empirical insights into corporeal awareness stem to a large extent from experimental designs, which allow temporarily altering the sense of ownership through multisensory stimulation (Botvinick & Cohen, 1998; H. H. Ehrsson, 2007; B. Lenggenhager, Tadi, Metzinger, & Blanke, 2007; Petkova & Ehrsson, 2008). Most famously, Botvinick and Cohen induced illusory ownership of an artificial hand by stroking a rubber hand placed in front of the participant in synchrony with the participant’s real hidden hand (Botvinick & Cohen, 1998). This phenomenological experience of ownership over the rubber hand is accompanied by objectively-measurable changes in a broad variety of processes ranging from basic physiological mechanisms (e.g., body temperature (Macauda et al., 2015; G. L. Moseley et al., 2008), nociception (Hansel, Lenggenhager, von Kanel, Curatolo, & Blanke, 2011; Romano, Pfeiffer, Maravita, & Blanke, 2014), cardiac signalling (Park et al., 2016), response to threat (H. Henrik Ehrsson, Wiech, Weiskopf, Dolan, & Passingham, 2007), and immunological responses (Barnsley et al., 2011)) up to high-level cognition (see e.g. (Maister, Slater, Sanchez-Vives, & Tsakiris, 2015) for a recent review). Empirical data thus suggest that the bodily self is highly plastic and based upon momentary sensory integration.

Next to our understanding of the nature of bodily self-consciousness and its disorders (Brugger & Lenggenhager, 2014), these data contribute towards developing methods to restore the bodily self where it is disturbed (G. L. Moseley, 2007; Pazzaglia, Haggard, Scivoletto, Molinari, & Lenggenhager, 2016). Strikingly however, most studies using multisensory stimulation in healthy participants or patients targeted the manipulation to bodily ownership of an additional and external, fake or virtual body (part), or more recently, two virtual representations of one’s own limb (Newport & Preston, 2011; Ratcliffe & Newport, 2017). In the above-described rubber hand illusion (RHI), the most striking phenomenological perception is not the feeling of disownership of the real hand but the feeling of ownership for the supernumerary rubber hand. In fact the two sensations are difficult to disentangle due to the nature of the RHI’s experimental design (Longo, Schüür, Kammers, Tsakiris, & Haggard, 2008). This “positive” embodiment stands in clear contrast to the clinical cases of disturbance of body ownership. While illusory ownership for a foreign body part has been described initially in certain cases of Somatoparaphrenia (Gerstmann, 1942), such neurological patients generally present with an altered sense of ownership for the own biological body. For example, in the more typical case of Somatoparaphrenia, patients lose the feeling of ownership for their own hand (Bottini, Bisiach, Sterzi, & Vallar, 2002). Similarly, Xenomelia patients feel like parts of their biological body do not belong to them (McGeoch et al., 2011). Even in the case of more general disturbances of the bodily self, for example the depersonalization syndrome, the own biological body does often not feel like being “self” anymore (Sierra, Baker, Medford, & David, 2005).

We here describe a new paradigm, the Real Hand Illusion (ReHI), designed to address this discrepancy between clinical reports and existing research paradigms in trying to alter the sense of ownership of one’s own biological limb in an Augmented Reality setup. During the illusion, participants view their own hand directly and from a first-person perspective through a head-mounted display (HMD) being touched either synchronously or asynchronously to the visual feedback (see Figure 1). The illusion was assessed with a questionnaire adapted from the classical RHI questionnaire (Botvinick & Cohen, 1998; Longo et al., 2008). As synchronous visuotactile stimulation has repeatedly been shown to increase perceived ownership, we predicted continued ownership during the synchronous stimulation and – more pertinent to the phenomenology described in clinical cases – decreased ownership of the own limb during the asynchronous condition. The latter could thus be regarded as temporarily mimicking the phenomenology found in Somatoparaphrenia or Xenomelia patients, i.e. the feeling of estrangement for their own limb.

**Figure 1.**
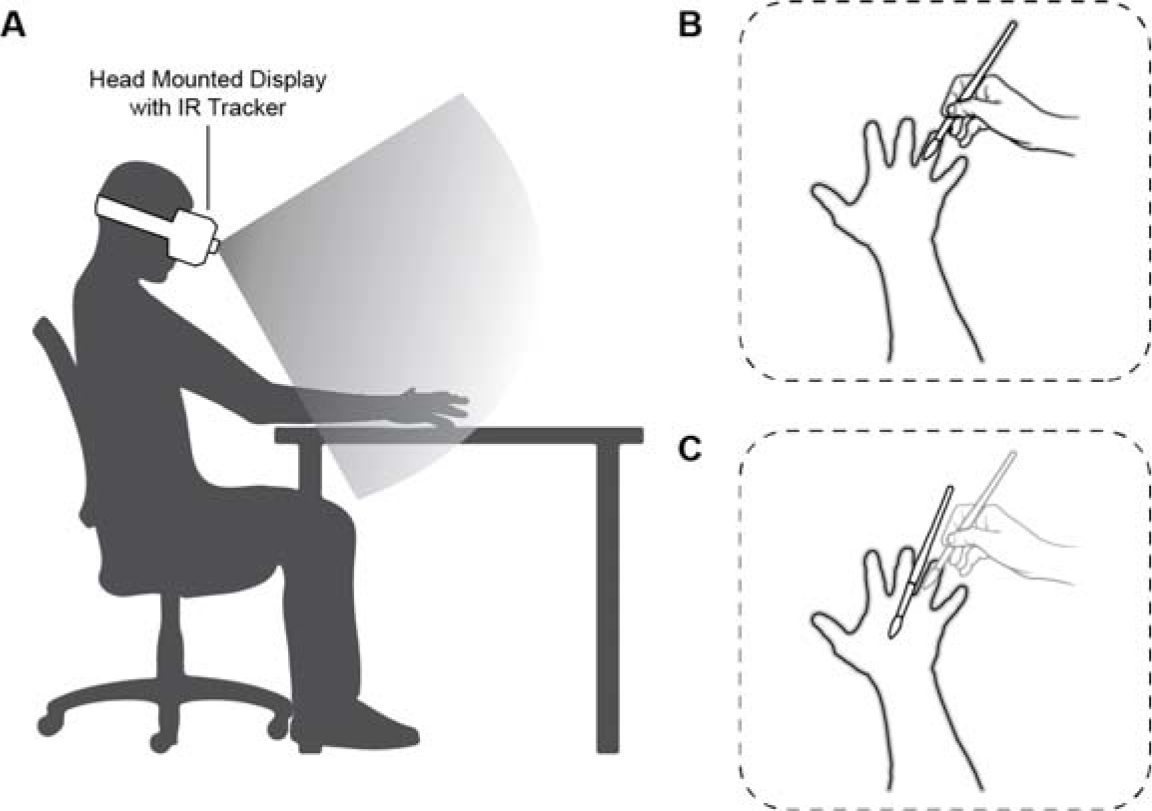
A. Participants are seated at a desk with their right arm resting on a pillow at their side. They wear the Head-Mounted Display with attached IR camera. The video feed of the camera is used to display the image of the participant’s own hand in the HMD. B. The biological and augmented limbs are aligned so that participants see their own hand in the correct anatomical position. In the control condition the feedback accurately presents the experimenter’s hand and the paintbrush providing synchronous (matching) visuotactile feedback. C. In the experimental condition, a 400ms delay is introduced in the visual feedback. Participants therefore feel the touch of the paintbrush (light grey) before seeing the paintbrush in the corresponding position.

Both Somatoparaphrenia and Xenomelia have been suggested to relate to alterations in multisensory bodily areas in the predominantly right-hemispheric posterior parietal areas (Hilti et al., 2013; McGeoch et al., 2011; Rode et al., 1992). As a consequence, both syndromes predominantly affect the left side of the body. In a second study, we thus investigated whether the feeling of disownership could be evoked more easily on the left as compared to the right hand in healthy participants. Based on previous literature on the rubber hand illusion which suggests stronger illusion for the left hand (Ocklenburg, Rüther, Peterburs, Pinnow, & Güntürkün, 2011), we hypothesized a stronger sensation of disownership during asynchronous stroking of the left as compared to the right hand.

In line with the idea of a right posterior parietal contribution to disorders of body ownership, we further investigated whether neuromodulation of parietal areas might alter the illusion in a systematic way. Brain imaging studies in individuals with Xenomelia have reported altered neural processes in the superior and inferior parietal lobe (Hilti et al., 2013; McGeoch et al., 2011; Oddo-Sommerfeld et al., 2018). Limb misidentification due to right-hemispheric damage has also been associated with parietal areas (Antoniello & Gottesman, 2017; Vallar & Ronchi, 2009). In line with this, neuromodulation through vestibular stimulation, which activates right parieto-insular areas (C. Lopez, Blanke, & Mast, 2012), has been shown to be helpful in Somatoparaphrenia (Rode et al., 1992) and consecutively also suggested as a therapeutic approach for Xenomelia ((Ramachandran & McGeoch, 2007); but see also (Bigna Lenggenhager, Hilti, Palla, Macauda, & Brugger, 2014)). Similarly, left anodal galvanic vestibular stimulation has been used to manipulate bodily ownership in healthy participants in a rubber hand illusion setup (Christophe Lopez, Lenggenhager, & Blanke, 2010). In a further, exploratory study, we used transcranial brain stimulation rather than indirect peripheral stimulation to alter activation of right parietal areas. We applied anodal and cathodal tDCS over the right superior parietal lobe normalised by a baseline sham stimulation. In line with previous literature we expected stimulation of right parietal networks to modulate body ownership.

## 2. MATERIALS AND METHODS

### 2.1 Setup

The technical setup follows the methods of our previous study (Bernal, Maes, & Kannape, 2016) as described in the following. A MacBook Pro Retina by Apple (Apple Inc., Cupertino, CA, USA), was used to render the visual feedback. The laptop had a dedicated AMD Radeon R9 M370X graphics card. The Oculus Rift DK2 (Subsidiary of Facebook, Menlo Park, CA, USA), Version 1.6 (SDK 0.5.0.1), was used to display the feedback. The HMD has a resolution of 960×1080pixels per eye, a horizontal field of view of 100°, and a refresh rate of 60Hz. Head orientation but not translation was tracked, as participants were asked to keep their head stationary during each trial. A LeapMotion controller (Leap Motion, Inc. San Francisco, CA, USA, Software Version 2.3.1) recorded the participant’s hand as well as the paintbrush, used to provide tactile feedback, using the integrated infra-red (IR) cameras. The visual stimuli used the resulting IR pass-through feed so that participants would see their own hand, as opposed to a rigged 3D model (cf. supplemental figure 1). Finally, the Unity game engine (Unity Technologies, San Francisco, CA, USA Version 5.1.3f) was used to render the stimuli in an otherwise empty virtual space.

### 2.2 Synchronous and Asynchronous Feedback

In order to change the delay of the visual feedback between the synchronous and asynchronous conditions, a buffer of the IR-feed was implemented using Leap’s Controller object within Unity. This maintains a frame history buffer of 60 frames. At 120fps sampling of the LeapMotion cameras, this provides up to half a second delay. Here, a 40-frame delay was used in order to produce a ~400ms delay during asynchronous feedback. This includes the intrinsic latency of the equipment which is as follows: tracking camera frame rate (120fps, ~8ms), tracking algorithm (4ms), display refresh rate (60Hz, ~17ms), and GPU calculations (~17ms) totalling to an intrinsic system delay of ~46ms (Bernal et al., 2016). Feedback in the synchronous condition was therefore achieved in under 50ms.

### 2.3 Tactile Feedback

Tactile feedback was provided using an ordinary flat, short-haired paint brush (size 10). The experimenter stroked the dorsum of the participant’s hand and fingers in different positions and directions for a total of three minutes. Unlike in previous limb ownership studies, only the participant’s hand was stroked, as opposed to an additional rubber hand. Accordingly, the visuo-tactile conflict in the asynchronous condition is purely temporal, and the visuo-tactile feedback in the synchronous condition exactly matches the actual stimulation. It should be noted that the experimental condition of the RHI, synchronous feedback, is in this case the control condition, Figure 1B; the asynchronous visuotactile stimulation, the control condition in the RHI, becomes the experimental condition, Figure 1C.

In studies 2 and 3, but not study 1, the three-minute illusion was preceded by one minute of synchronous, visual only, feedback. Here, participants were asked to move their hands in order to familiarise themselves with the environment, get accustomed to the feedback, and appreciate that they have full control of their own arm in the VR space.

### 2.4 ReHI Questionnaire

Following each trial, participants were asked to write down any comments they had about their perception of the illusion (open feedback). Phenomenological aspects of the illusion were then systematically assessed with a questionnaire adapted from the classical RHI questionnaire (Botvinick & Cohen, 1998; Longo et al., 2008). Questions one through six were scored positively from 0 to 10, whereas questions seven through ten pertain to the feeling of disownership and were therefore reverse-coded for the analysis (0-> 10, 1 ->9, etc.). This was done so that all ten questions were combined to calculate an overall illusion-score with a possible range of 0 to 100. A score of 100 represents the highest possible ‘embodiment score’, whereas a score of 0 would reflect a complete loss of ownership and agency of the seen hand.

### 2.5 Participants

Study 1 (N=20, age µ=21±1years, 12 female) investigated the illusion based on the individual questions and an overall score (see figure 2 for boxplots). A t-test comparing the illusion score between the two conditions of interest (i.e., synchronous versus asynchronous stroking) revealed a significant effect (t(19)=4.58, p<0.001, Cohen’s dz = 1.02) between conditions. This Cohen’s d was used for power calculations in studies 2 and 3. Participants for all three studies were right-handed (self-reported) and had normal or corrected to normal vision.

**Figure 2.**
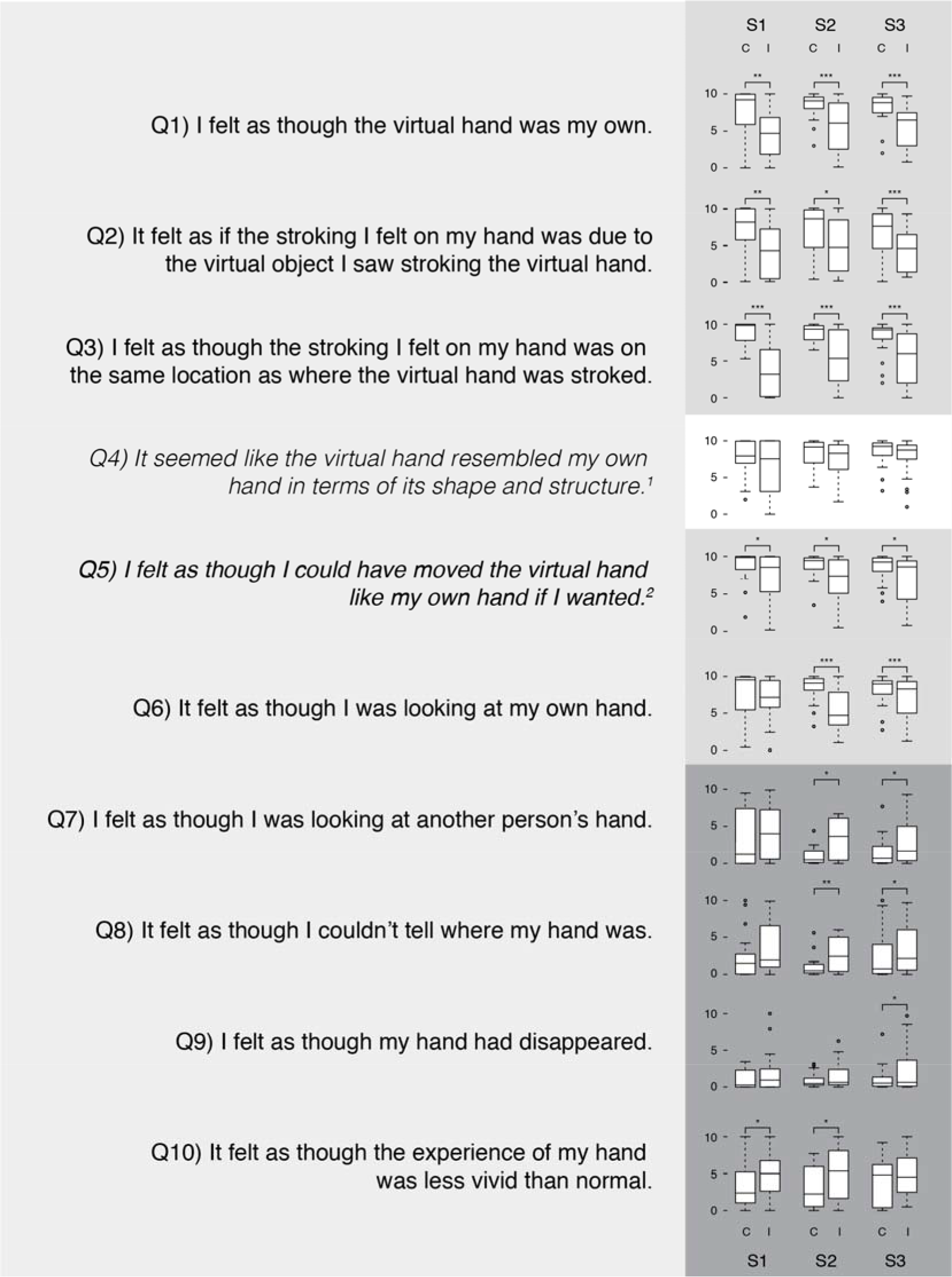
Questions and box plot with interquartile ranges of the results across the three studies – Questions 1 through 6 were adapted from previous studies on the RHI. Question 4 is a control question and question 5 addresses participants’ sense of agency. Questions 7 through 10 were included to directly address disownership aspects of the ReHI. Participants in all three cohorts rated embodiment higher given synchronous visuotactile feedback about their upper limb in the control condition compared to the asynchronous feedback during the ReHI. All asterisks indicate NHST significance. Data for S2 and S3 (sham stimulation) are taken from left hand.

A power calculation (G*Power (Faul, Erdfelder, Lang, & Buchner, 2007)) indicated that a sample size of 15 would be required to detect an effect size of dz = 1.02, with the alpha level set at .05, with power of .95 (two-tailed). Twenty participants were recruited for study 2 (age: µ=21.55±2.48years, 10 females).

Power calculation for the exploratory neurostimulation study was further informed by Kammers and colleagues effect size of approximately *d* = 0.6, reported in their rTMS study on the RHI (Marjolein P. M. Kammers et al., 2009). Using a paired samples t-test to contrast two stimulation conditions, to reach 80% power with this effect size (alpha level = .05) would require 24 participants. Twenty-six participants (age µ=21.32±8.31years, 16 male) were recruited and completed the tDCS paradigm (study 3). All participants refrained from consuming caffeine for at least three hours prior to the tDCS stimulation. All studies had been approved by the University of Central Lancashire’s Ethical committee (Protocol PSYSOC336).

### 2.6 Transcranial direct current stimulation

TDCS over the right SPL was used to change cortical excitability (see Figure 3). In one condition, anodal stimulation was applied over the right SPL with the aim of increasing cortical excitability, with the cathodal electrode as the reference. In another condition, cathodal stimulation was applied over the right SPL with the aim of decreasing cortical excitability. The third condition was the sham stimulation condition, which acted as a baseline against which to compare active stimulation. Participants attended two separate sessions, separated by at least three days. Participants always completed the anodal and cathodal stimulation conditions in separate sessions, to avoid after-effects from one condition affecting another condition. Half the participants additionally completed the sham condition at the start of the first session, while the remaining half completed the sham condition at the start of the second session. The stimulation was applied double-blinded using pre-determined codes stored on the tDCS equipment, which determined the type of stimulation applied (anodal, cathodal or sham) without the experimenter’s knowledge.

**Figure 3.**
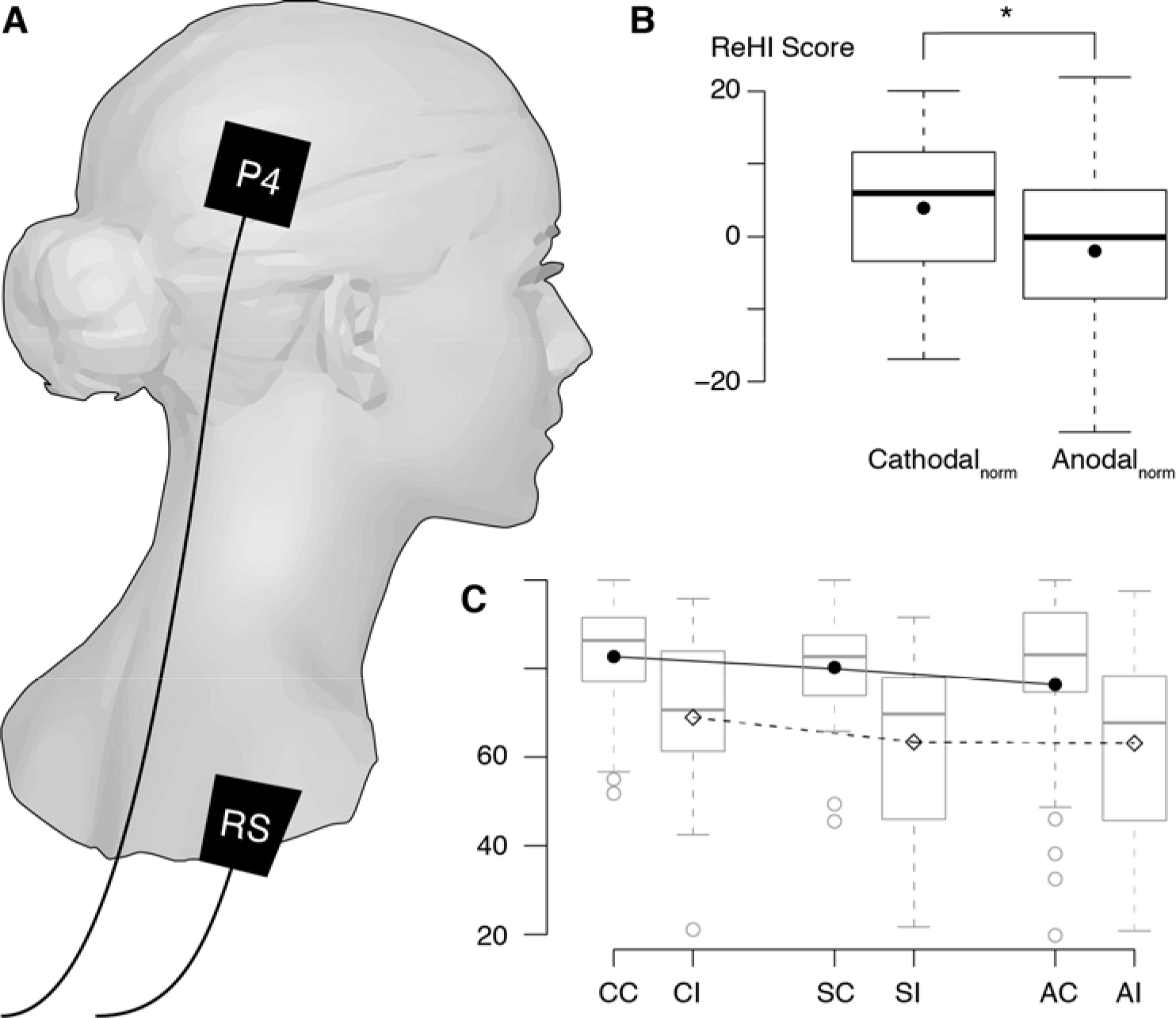
Effects of Transcranial Direct Current Stimulation on ReHI Strength. A Electrodes were placed on the P4 position based of the international 10-20 system and the right shoulder. B The main contrast indicates that anodal stimulation over P4 significantly decreased embodiment across both conditions compared to cathodal stimulation. (SHAM scores were subtracted for baseline correction.) C Breakdown of questionnaire scores across control and ReHI conditions and stimulation type. All boxplots indicate medians and Interquartile ranges; means are indicated by solid circles or diamonds (CC/CI = Cathodal Control/Illusion; SC/SI = Sham Control/Illusion; AC/AI = Anodal Control/Illusion).

The electrodes were positioned with the aid of an EASYCAP 21 EEG cap (EASYCAP, Herrsching), with the scalp electrode positioned over the P4 region according to the international 10-20 system. The P4 electrode is located approximately over the right superior parietal lobe (Herwig, Satrapi, & Schönfeldt-Lecuona, 2003) and has previously been used to target this region (Ono, Mikami, Fukuyama, & Mima, 2016). The reference electrode was positioned over the ipsilateral shoulder, held in place with a rubber strap (cf. Figure 3).

In the two tDCS conditions, participants received 1200s (including 8s of fade-in and 8s of fade-out) of tDCS, using a NeuroConn DC-Stimulator Plus (NeuroConn, Germany). A 1.5mA current was delivered through 25cm^2^ saline-soaked sponges (0.9% NaCl solution), held in place on the participant’s scalp by rubber straps (current density of 0.06mA/cm^2^). Stimulation was applied for 600s before task onset, and continued 600s after task onset.

Sham stimulation consisted of stimulation applied for 38s, before dropping to regular pulses of 115µA (lasting 3ms) every 550ms, which gives an average current strength of 0.002mA. This level of stimulation is far lower than required to cause changes in cortical excitability but allows monitoring of impedance (which could indicate poor electrode contact or disconnection).

### 2.7 Analysis and Data Availability

Paired frequentist and Bayesian t-tests were conducted in JASP (JASP Team, 2018). These tests were two-tailed for NHST and directional for calculation of BF due to clear directional hypothesis of the illusion effect. Significance thresholds were set to p<.05 and BF>3 (BF<.3) respectively. NHST was used in combination with power-calculations; Bayesian statistics (Dienes, 2014) are included as potential evidence in favour of the null. Cronbach’s alpha was calculated in R using R-Studio (R Development Core Team, 2017) and the Psych package (Revelle, 2017). All data will be made available to the readers via a public Open Science Framework project (http://osf.io/wbp59).

## 3. RESULTS

As this is a novel illusion, we first set out to investigate the effects of the Real Hand Illusion on the phenomenology of the bodily self, focusing on hand ownership (Q1), location of touch (Q2-3), agency (Q5), and aspects of disownership (Q7-Q10). To compare the overall illusion across conditions, a combined score was created based on all questions; for this, disownership questions were reverse-coded.

### 3.1 Study 1 – Limb Disownership

In study 1, twenty participants completed the synchronous and asynchronous feedback conditions and the ten-item questionnaire. As illustrated in Figure 2, participants rated the seen hand to feel less like their own during the illusion (Q1 control: 7.58±3.21 to illusion: 4.46±3.21 (µ±σ), p=.002, BF=36.50). Similarly, they reported that it felt less likely that the stroking they felt on their hand was due to the stroking on the seen hand (Q2: 7.06±3.44 to 4.16±3.46, p=.009, BF=10.96), and less likely that it was stroked in the same location (Q3: 8.87±1.47 to 3.85±3.51, p<.001, BF=2425.24). Questions 4 and 6 are based on control questions from the RHI questionnaire but may reflect aspects of limb disownership addressed in the current illusion. In study 1, participants did not perceive the shape and structure of the hand to differ between conditions (p=.051, BF=2.66), nor did they feel that it felt less like looking at their own hand when receiving asynchronous feedback (Q6: p=.510, BF=.28). However, participants did report an effect of the ReHI on their (hypothetical) ability to move the seen hand (“feeling of agency” Q5: 8.81±2.11 to 7.16±3.37, p=.021, BF=5.41). With respect to the disownership questions, participants reported that their hands felt less vivid than normal during the illusion (Q10: 6.83±2.88 to 5.15±2.95, p=.016, BF=6.89). Questions Q7-9 were not significantly different between conditions in study 1 (all p>.085, all BF inconclusive .45<BF<.1.75).

All questions were combined to calculate an overall illusion score; disownership questions were reverse-coded. This led to a possible range of scores from 0 to 100, based on the six positively and 4 reverse-coded questions. As hypothesised, participants rated the Real Hand Illusion questions significantly more positively in the control condition (77.53±13.08) than during the illusion (58.47±17.27, p<.001, BF=292.66) designed to induce a loss of ownership.

### 3.2 Study 2 – Lateralisation

In study 2, the lateralization of body representation and its malleability were investigated. It was hypothesised that the strength of the illusion would be higher for participants’ left hand compared to their right. A 2 x 2 repeated measures ANOVA, based on the overall illusion strength, confirmed our data from study 1 with respect to the main effect of the illusion. The total score significantly dropped from 83.95±13.10 during synchronous feedback to 66.36±20.06 during the asynchronous feedback of the illusion (main effect of illusion F(1,19)=16.04, p<.001, η2=.46). There was no overall effect of laterality (p=.107); however, there was a significant interaction between the two factors (F(1,19)=4.84, p=.026, η2=.24): Questionnaire scores were lower for the left than the right hand during the illusion but not in the control condition. As hypothesised, the illusion was stronger for the left hand (paired t-test: µ-difference: 6.41±2.69, t(19)=2.38, p=.028). The ANOVA results were corroborated in Bayesian t-tests. The data strongly support a main effect of illusion (C>I: BF=92.98) but not of lateralization (R≠L, BF=.78). The interaction, indicated by the left-right differences in both conditions, was also supported by the data (RI-LI > RC-LC: BF=4.61), resulting from the lateralisation difference during the illusion (LI < RI: BF=4.43).

Inspecting the individual scores for the left hand only (for comparison with the study 3) confirmed significant effects on the ‘classical’ ownership questions (Q1-Q3) as in study 1 (all p<.013, all BF>7.95). Similarly, as in study 1, the ReHI affected the perceived ability to move (Q5: p=.012, BF=8.74) and caused the own hand to feel less vivid during the illusion (Q10: p=.038, BF=3.33). In addition, participants in study 2 reported that it felt less like they were looking at their own hand (Q6: p<.001, BF=744.42) but more likely that they were looking at somebody else’s hand (Q7: p=.001, BF=53.29). Finally, participants felt more strongly that they could not tell where their hand was during the illusion (Q8: p=.005, BF=18.74).

### 3.3 Study 3 – Neurostimulation

In study 3, we set out to investigate the involvement of right parietal networks in maintaining body representation during the Real Hand Illusion. In two separate sessions, participants completed the ReHI while receiving sham, anodal, or cathodal tDCS stimulation, with the scalp electrode placed over the right superior parietal lobe. As this was an exploratory study, we were mainly interested in the overall effect of stimulation on ReHI score and the main contrast of anodal versus cathodal stimulation. In order to normalise the data, we therefore subtracted the average ReHI score during sham stimulation from anodal and cathodal scores. This resulted in a 2 x 2 repeated measures design with factors Illusion (synchronous, asynchronous) and Stimulation (anodal, cathodal). In-line with the studies 1 and 2, we report a significant main effect of the illusion in a third, independent participant pool (F(1,25)=25.54, p<.001, η2=.51, BF_C>I_=1474.21). We additionally observed a main effect of Stimulation with ReHI scores being higher during cathodal stimulation than during anodal stimulation (F(1,25)=5.35, p=.029, η2=.18, BF_C>A_=3.81, BF_C≠A_=1.94, see Figure 3b), although there was no interaction between illusion condition and stimulation condition (F(1, 25) = 0.02, p = .89).

Figure 2 illustrates the box-plots for the ReHI score in the sham condition (left hand). Akin to both the studies 1 and 2, participants rated ownership questions Q1-3 significantly lower during the illusion (all p<.001, all BF>77.55). Questions regarding perceived movement (Q5), and ownership/disownership questions Q6-9 were scored also significantly lower during the ReHI (all p<.025, all BF>4.39). Only Q10, regarding the perceived vividness of one’s own hand did not reach significance in this condition (p=.077, BF=1.72).

### 3.4. Questionnaire Reliability

As this is a novel illusion accompanied by a newly designed questionnaire, we ran a reliability analysis across the three data sets (N=66) – once for the control condition, and once for the illusion condition. Cronbach’s alpha for the questionnaire responses in the control condition was α_control_=0.79, in the illusion α_illusion_=0.85, indicating good internal consistency across the three cohorts.

## 4. DISCUSSION

We here introduce a body-illusion that, for the first time, directly reduces the feeling of ownership over one’s biological limb, without relying on feedback from a supernumerary proxy such as a rubber hand. Participants view their own arm, in Augmented Reality, from an embodied, first person perspective but receive feedback that contains a temporal, visuotactile conflict. In a series of three studies, in three separate participant pools, we demonstrate that the asynchronous feedback in the Real Hand Illusion causes participants to rate ownership (and the sense of agency) over their own hand and related sensations significantly lower than in the synchronous control condition. In an open feedback, multiple participants independently described the phenomenology of the illusion akin to “pins and needles”, the tickling sensation that often occurs after transient paraesthesia.

The Real Hand Illusion, which could be considered as an inverse of the classical rubber hand illusion, hence modulates body ownership by introducing a controlled mismatch into bottom-up multisensory integration. Our data suggest that this weakened embodiment is pronounced for the left hand, supporting the right hemispheric dominance hypothesis of body representation as reflected in neuropsychological case reports and previous ownership illusion studies in healthy participants. Transcranial direct current stimulation over the right superior lobule modulated the overall strength of limb ownership, corroborating the role of right posterior parietal networks for multisensory representations of the body and self.

### 4.1 Phenomenology

Embodiment, ownership, and the sense of agency have been argued to be matters of “a very thin phenomenal awareness” (Gallagher, 2007) and to only form the “background of mental life” (Longo et al., 2008). Often, we only fully become aware of these processes when they break down – which can have severe consequences (a collection of accounts are available in 54, 55). While research into the sense of agency has managed to address this by experimentally modulating a loss of control (48–51)(Franck et al., 2001; Kannape & Blanke, 2013; Leube et al., 2003; Nielsen, 1963), a symptom that is often found in clinical conditions (see e.g. (Blakemore, Wolpert, & Frith, 2002)), ownership studies have instead investigated the opposite: a ‘positive’ ownership of an artificial limb such as a rubber hand or foot (Bigna Lenggenhager, Hilti, & Brugger, 2015), or even two versions of one’s own hand (Ratcliffe & Newport, 2017) (an interesting conundrum of this study is that the ‘fake’ hand is actually based on the participant’s own hand, leading to the question of whether one can disown one representation of one’s hand over another). Analog to sensorimotor studies delineating the spatiotemporal limits of agency, we have here introduced a paradigm to directly induce disownership of one’s own limb: where participants report a loss of control over their actions in agency research, participants in the ReHI perceive a weakened ownership over their own hand and arm.

The topic of disownership is a somewhat contentious area with respect to the RHI. There is a general consensus that the rubber hand is embodied during synchronous feedback (see e.g. (Serino et al., 2013)) – arguably only into the body image (for perception) but not the body schema (for action) (M.P.M. Kammers, de Vignemont, Verhagen, & Dijkerman, 2009). However, this does not imply that one’s own limb is simultaneously disembodied. While arguments have been made for disembodiment, either by asking directly about disownership of the real hand (Preston, 2013) or by inferring from physiological data (drop in skin temperature (G. L. Moseley et al., 2008; Salomon, Lim, Pfeiffer, Gassert, & Blanke, 2013) or alterations in immune response (Barnsley et al., 2011), but see (de Haan et al., 2017) for a critical account) previous studies have relied on including a supernumerary limb, and evidence suggests that multiple representations of the hand might co-exist (H. Henrik Ehrsson, 2009; McGonigle et al., 2002). Ultimately, the strongest statement to be made in favour of disembodiment from these previous paradigms is that the supernumerary hand replaces or “functionally suppresses” the participant’s actual hand (Longo et al., 2008).

Importantly, the phenomenology of the ReHI extends previous findings as it precludes both supernumerary embodiment and the replacement of the actual limb representation. There is only one arm. Rather than being indicative of a malleability to multisensory body illusions or an ability to incorporate supernumerary limbs, the current findings hence suggest that limb representation can directly be attenuated, without “tricking the brain” by including a proxy. It is therefore not so much illustrating the malleability of limb representation but questions its actual permanence: contradicting William James famous premise (James, 1890), the body may not always be there.

### 4.2 Handedness and Lateralisation

Study 2 illustrates that the phenomenological experience of the ReHI was stronger for the left hand as opposed to the right. This difference in lateralisation was specific to the illusion condition as no lateralisation was observed in the control condition. The findings are in line with mounting evidence that Xenomelia, similar to Somatoparaphrenia, more often than not affects the left side of the body (Brugger & Lenggenhager, 2014; McGeoch et al., 2011). While an argument is to be made that lateralisation may be a result of handedness, and stronger bodily illusions have been reported for the non-dominant hand (Brugger & Meier, 2015), evidence from the RHI suggests there is a right-hemispheric dominance for sense of body ownership independent of handedness ((Ocklenburg et al., 2011), but also see (Smit, Kooistra, van der Ham, & Dijkerman, 2017)). Taken together, this suggests that the ReHI is mediated by multisensory bodily areas in right-hemispheric posterior parietal areas (Hilti et al., 2013; McGeoch et al., 2011; Rode et al., 1992), further motivating the exploratory neurostimulation study to focus on the left upper-limb.

### 4.3 Neurostimulation modulates Limb Disownership

The results from Study 3 indicated that application of tDCS over the right SPL modulated the experience of ownership, dependent on the polarity of stimulation. Specifically, anodal stimulation led to reduced feelings of ownership over the limb, while cathodal stimulation increased feelings of ownership. It should be noted that this was an exploratory study, aiming to link the ReHI to activity in the parietal cortex; future studies should therefore aim to test under which conditions the effect of stimulation holds. Our data suggests that stimulation affected the experience of ownership during both the illusion and control conditions, suggestive of a broad effect of stimulation that is not dependent on synchronous visuotactile feedback. Previous studies have shown that transcranial magnetic stimulation (TMS) applied to the right temporoparietal junction (TPJ) disrupts the rubber hand illusion for body-like objects (but not other objects) (Tsakiris, Costantini, & Haggard, 2008). Clinically, low-frequency repetitive TMS applied to the TPJ may decrease the frequency of auditory hallucinations in patients with a diagnosis of schizophrenia, a symptom frequently linked to loss of agency or ownership over self-generated speech (P. Moseley, Fernyhough, & Ellison, 2013; Slotema, Blom, van Lutterveld, Hoek, & Sommer, 2014). Taken together, these findings suggest that non-invasive neurostimulation is capable of affecting the perception of bodily ownership and support the therapeutic potential of neurostimulation in disorders of bodily ownership.

### 4.4 ReHI Considerations

In contrast to the on-going, artificially synchronised visuotactile feedback required by the RHI, the ReHI relies on exploiting the same bottom-up multisensory integration processes by introducing a temporal mismatch to disrupt body ownership. In addition to the aforementioned conceptual advantages, this has a number of practical advantages. One, the sensation of the mismatch is immediate. Unlike the RHI, which relies on continued synchronous feedback from two distinct, visuotactile sources and has reported onset times between 10-50 seconds (H. H. Ehrsson, 2004; Kalckert & Ehrsson, 2017) the ReHI relies on a hard-coded temporal mismatch from a single source. Two, the multisensory mismatch is unresolvable, making the illusion very stable. Participants feel the touch on the back of their hand before receiving visual feedback. As the position of the subsequent touch-location is unpredictable, there cannot be an adaptation to the conflicting sensory information. Three, the mismatch is purely temporal. Whereas inadvertent spatiotemporal incongruences occur in both synchronous and asynchronous conditions in the RHI, only a single hand is stimulated in the ReHI. Four, and continuing this point, the control condition is very accurate, as the perceived location of the touch exactly corresponds to the seen location. The technical setup precludes an unwanted (spatial) mismatch, apart from the intrinsic (temporal) delay. Finally, the setup is portable and easily implemented, making it a promising tool for clinical studies and potential outpatient treatment. The paradigm relies on a simple (video) buffer, making it adaptable to a range of mobile devices and commercially available research platforms.

### 4.5 Limitations and Future Directions

As this is a novel paradigm, there are a number of open questions to address. For example, we based our questionnaire on widely used RHI items but included four new questions directly addressing limb disownership (Q7-10). A follow-up study that investigates an expanded questionnaire using principal component analysis, similar to the approach taken by Longo and colleagues (Longo et al., 2008) may be able to improve the reporting of the phenomenology of the ReHI. Linking back to the “pins and needles” sensation reported by participants, physiological reactions to the illusion should be investigated following previous protocols on the RHI (body temperature (Macauda et al., 2015; G. L. Moseley et al., 2008), nociception (Hansel et al., 2011; Romano et al., 2014), cardiac signalling (Park et al., 2016), response to threat (H. Henrik Ehrsson et al., 2007), and immunological responses (Barnsley et al., 2011)) up to high-level cognition (Maister et al., 2015)).

A further line of inquiry should address the clinical aspects of limb disownership by working with patient populations. Individuals with Xenomelia show an enhanced response for the affected limb in a rubber hand illusion type of setup (Bigna Lenggenhager et al., 2015). Does the same hold true for the ReHI and how does its phenomenology compare to the sensation of “over-completeness” described by these individuals? Does it capture aspects of loss of limb ownership experienced in Somatoparaphrenia? Applying the ReHI in individuals with Xenomelia may recreate the reported feeling of disownership – or, by inducing disownership over an unwanted limb – create a cessation in the dysphoric feeling of over-completeness. Similar to the pretend behaviour exercised by these individuals (First, 2005; L. Fischer, 2015), this may offer a temporary relief, if not a treatment. Similarly, research with such a cohort will be relevant to understanding permanence of body representation. If the loss of ownership is a gradual process, it could potentially be tracked longitudinally using the ReHI and further related to the frequency and duration of pretend behaviour over time. The exploratory tDCS results further merit investigation of neurostimulation as a therapeutic possibility (although this may evoke ethical questions pertaining to the identity of individuals with Xenomelia). Finally, applying an analogue paradigm to the full body (Blanke, Slater, & Serino, 2015) may provide an experimental link to investigating aspects of depersonalisation in the general population (Sierra & David, 2011).

## 5. CONCLUSION

The Real Hand Illusion introduced here offers a direct way of modulating limb ownership in healthy individuals. It does so without relying on a proxy, but by introducing a temporal, visuotactile mismatch into bottom-up processed feedback about one’s own limb in Augmented Reality. These findings are corroborated by two additional studies in independent participant pools linking the illusion to right posterior parietal networks for multisensory representations of the body and self. By directly investigating the loss of ownership of one’s own limb, analogue to research into the sense of agency, the Real Hand Illusion opens up the possibility of more adequately addressing the majority of clinical cases of altered body ownership; further, it provides a novel method of investigating body representation and its permanence in healthy individuals.

## SUPPLEMENTAL FIGURE

**Supplemental Figure 1.**
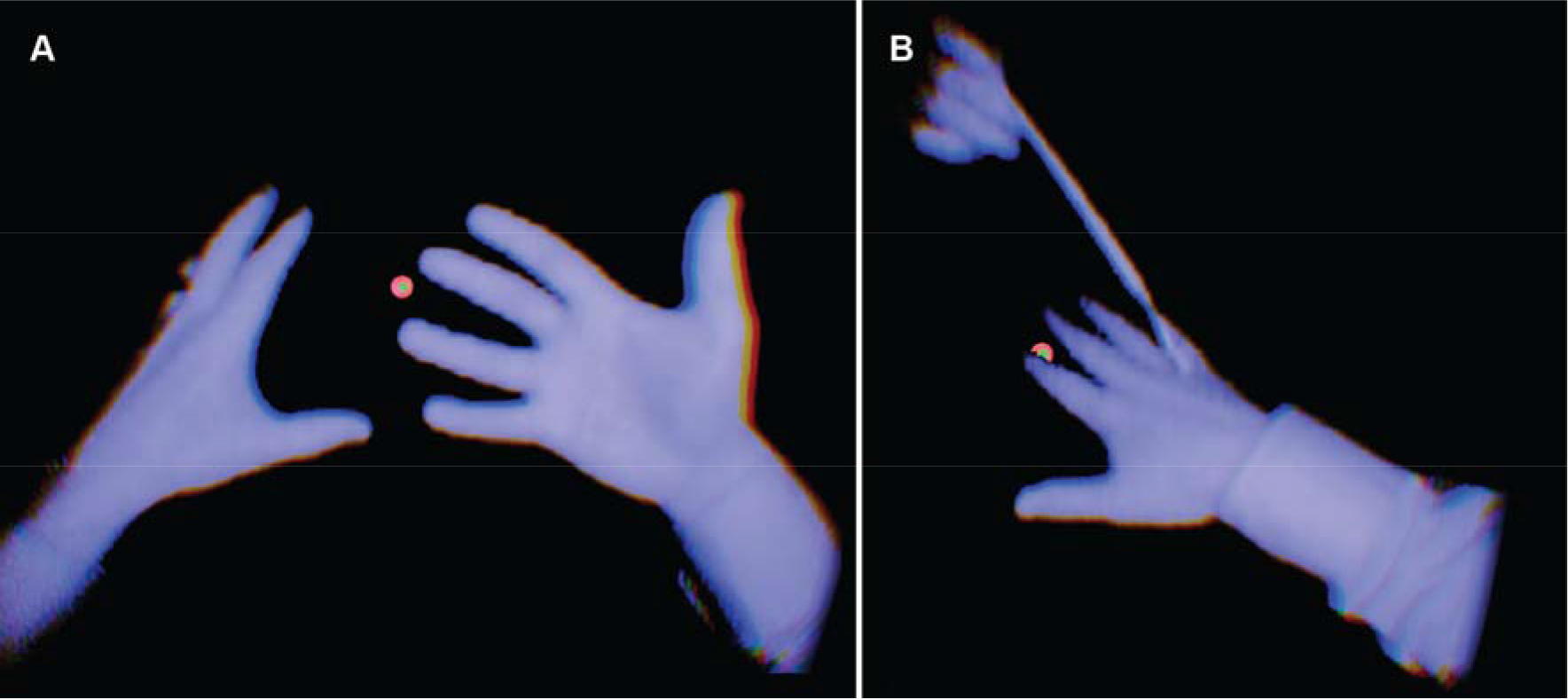
Visual Feedback during the ReHI. ***A.*** Participants viewed an Infra-Red feed of their own hands, from a first-person perspective, in an otherwise empty virtual environment (cropped from left eye view). Before either condition, participants were provided with real-time feedback and asked to move their hands around the virtual space. ***B.*** During the control condition (real-time) and the ReHI condition (400ms visual delay), participants received visuo-tactile stimulation via a simple paintbrush. Perceiving temporally conflicting information about their own limb, led participants to report a loss of ownership of their own limb, compared to the control condition.

**FUNDING** BL was supported by the Swiss National Science Foundation (grant number 170511).

**AUTHOR CONTRIBUTIONS** Original Paradigm: OK, ET; Experimental Design: all authors; Data Collection: ET, OK; Data Analysis: all authors; Manuscript Preparation: all authors

## REFERENCES

Antoniello, D., & Gottesman, R. (2017). Limb Misidentification: A Clinical-Anatomical Prospective Study. The Journal of Neuropsychiatry and Clinical Neurosciences, 29(3), 284–288. https://doi.org/10.1176/appi.neuropsych.16090169

Barnsley, N., McAuley, J. H., Mohan, R., Dey, A., Thomas, P., & Moseley, G. L. (2011). The rubber hand illusion increases histamine reactivity in the real arm. Current Biology, 21(23), R945–R946. https://doi.org/10.1016/j.cub.2011.10.039

Bernal, G., Maes, P., & Kannape, O. A. (2016). The temporal limits of agency for reaching movements in augmented virtuality (pp. 002896–002899). IEEE. https://doi.org/10.1109/SMC.2016.7844679

Blakemore, S. J., Wolpert, D. M., & Frith, C. D. (2002). Abnormalities in the awareness of action. Trends Cogn Sci, 6, 237–242.

Blanke, O. (2012). Multisensory brain mechanisms of bodily self-consciousness. Nature Reviews. Neuroscience, 13(8), 556–571. https://doi.org/10.1038/nrn3292

Blanke, O., Slater, M., & Serino, A. (2015). Behavioral, Neural, and Computational Principles of Bodily Self-Consciousness. Neuron, 88(1), 145–166. https://doi.org/10.1016/j.neuron.2015.09.029

Bottini, G., Bisiach, E., Sterzi, R., & Vallar, G. (2002). Feeling touches in someone else’s hand. Neuroreport, 13(2), 249–252.

Botvinick, M., & Cohen, J. (1998). Rubber hands “feel” touch that eyes see. Nature, 391, 756.

Brugger, P., & Lenggenhager, B. (2014). The bodily self and its disorders: neurological, psychological and social aspects. Current Opinion in Neurology, 27(6), 644–652. https://doi.org/10.1097/WCO.0000000000000151

Brugger, P., & Meier, R. (2015). A New Illusion at Your Elbow. Perception, 44(2), 219–221. https://doi.org/10.1068/p7910

de Haan, A. M., Van Stralen, H. E., Smit, M., Keizer, A., Van der Stigchel, S., & Dijkerman, H. C. (2017). No consistent cooling of the real hand in the rubber hand illusion. Acta Psychologica, 179, 68–77. https://doi.org/10.1016/j.actpsy.2017.07.003

Descartes, R., & Cottingham, J. (2013). Meditations on first philosophy: with selections from the objections and replies◻; a Latin-English edition. Cambridge, England: Cambridge University Pr.

Dienes, Z. (2014). Using Bayes to get the most out of non-significant results. Frontiers in Psychology, 5. https://doi.org/10.3389/fpsyg.2014.00781

Ehrsson, H. H. (2004). That’s My Hand! Activity in Premotor Cortex Reflects Feeling of Ownership of a Limb. Science, 305(5685), 875–877. https://doi.org/10.1126/science.1097011

Ehrsson, H. H. (2007). The experimental induction of out-of-body experiences. Science, 317, 1048.

Ehrsson, H. Henrik. (2009). How many arms make a pair? Perceptual illusion of having an additional limb. Perception, 38(2), 310–312. https://doi.org/10.1068/p6304

Ehrsson, H. Henrik, Wiech, K., Weiskopf, N., Dolan, R. J., & Passingham, R. E. (2007). Threatening a rubber hand that you feel is yours elicits a cortical anxiety response. Proceedings of the National Academy of Sciences of the United States of America, 104(23), 9828–9833. https://doi.org/10.1073/pnas.0610011104

Faul, F., Erdfelder, E., Lang, A.-G., & Buchner, A. (2007). G*Power 3: A flexible statistical power analysis program for the social, behavioral, and biomedical sciences. Behavior Research Methods, 39(2), 175–191. https://doi.org/10.3758/BF03193146

First, M. B. (2005). Desire for amputation of a limb: paraphilia, psychosis, or a new type of identity disorder. Psychological Medicine, 35(6), 919–928.

Franck, N., Farrer, C., Georgieff, N., Marie-Cardine, M., Dalery, J., d’Amato, T., & Jeannerod, M. (2001). Defective recognition of one’s own actions in patients with schizophrenia. Am J Psychiatry, 158, 454–459.

Gallagher, S. (2007). The Natural Philosophy of Agency. Philosophy Compass, 2, 347–357.

Gerstmann, J. (1942). PROBLEM OF IMPERCEPTION OF DISEASE AND OF IMPAIRED BODY TERRITORIES WITH ORGANIC LESIONS: RELATION TO BODY SCHEME AND ITS DISORDERS. Archives of Neurology & Psychiatry, 48(6), 890. https://doi.org/10.1001/archneurpsyc.1942.02290120042003

Hansel, A., Lenggenhager, B., von Kanel, R., Curatolo, M., & Blanke, O. (2011). Seeing and identifying with a virtual body decreases pain perception. Eur J Pain, 15, 874–879. https://doi.org/10.1016/j.ejpain.2011.03.013

Herwig, U., Satrapi, P., & Schönfeldt-Lecuona, C. (2003). Using the international 10-20 EEG system for positioning of transcranial magnetic stimulation. Brain Topography, 16(2), 95–99.

Hilti, L. M., Hänggi, J., Vitacco, D. A., Kraemer, B., Palla, A., Luechinger, R., … Brugger, P. (2013). The desire for healthy limb amputation: structural brain correlates and clinical features of xenomelia. Brain: A Journal of Neurology, 136(Pt 1), 318–329. https://doi.org/10.1093/brain/aws316

Husserl, E. (2002). Ideen zu einer reinen Phänomenologie und phänomenologischen Philosophie: allgemeine Einführung in die reine Phänomenologie (6. Aufl., unveränd. Nachdr. der 2. Aufl. 1922). Tübingen: Niemeyer.

James, W. (1890). The principles of psychology. New York, NY: Dover. JASP Team. (2018). JASP (Version 0.8.5). Retrieved from https://jasp-stats.org/

Kalckert, A., & Ehrsson, H. H. (2017). The Onset Time of the Ownership Sensation in the Moving Rubber Hand Illusion. Frontiers in Psychology, 8. https://doi.org/10.3389/fpsyg.2017.00344

Kammers, Marjolein P. M., Verhagen, L., Dijkerman, H. C., Hogendoorn, H., De Vignemont, F., & Schutter, D. J. L. G. (2009). Is This Hand for Real? Attenuation of the Rubber Hand Illusion by Transcranial Magnetic Stimulation over the Inferior Parietal Lobule. Journal of Cognitive Neuroscience, 21(7), 1311–1320. https://doi.org/10.1162/jocn.2009.21095

Kammers, M.P.M., de Vignemont, F., Verhagen, L., & Dijkerman, H. C. (2009). The rubber hand illusion in action. Neuropsychologia, 47(1), 204–211. https://doi.org/10.1016/j.neuropsychologia.2008.07.028

Kannape, O. A., & Blanke, O. (2013). Self in motion: sensorimotor and cognitive mechanisms in gait agency. Journal of Neurophysiology, 110(8), 1837–1847. https://doi.org/10.1152/jn.01042.2012

L. Fischer, M. (2015). Body Integrity Identity Disorder: Development and Evaluation of an Inventory for the Assessment of the Severity. American Journal of Applied Psychology, 4(3), 76. https://doi.org/10.11648/j.ajap.20150403.15

Lenggenhager, B., Tadi, T., Metzinger, T., & Blanke, O. (2007). Video ergo sum: manipulating bodily self-consciousness. Science, 317, 1096–1099.

Lenggenhager, Bigna, Hilti, L., & Brugger, P. (2015). Disturbed body integrity and the “rubber foot illusion.” Neuropsychology, 29(2), 205–211. https://doi.org/10.1037/neu0000143

Lenggenhager, Bigna, Hilti, L., Palla, A., Macauda, G., & Brugger, P. (2014). Vestibular stimulation does not diminish the desire for amputation. Cortex; a Journal Devoted to the Study of the Nervous System and Behavior, 54, 210–212. https://doi.org/10.1016/j.cortex.2014.02.004

Leube, D. T., Knoblich, G., Erb, M., Grodd, W., Bartels, M., & Kircher, T. T. (2003). The neural correlates of perceiving one’s own movements. Neuroimage, 20, 2084–2090.

Longo, M. R., Schüür, F., Kammers, M. P. M., Tsakiris, M., & Haggard, P. (2008). What is embodiment? A psychometric approach. Cognition, 107(3), 978–998. https://doi.org/10.1016/j.cognition.2007.12.004

Lopez, C., Blanke, O., & Mast, F. W. (2012). The human vestibular cortex revealed by coordinate-based activation likelihood estimation meta-analysis. Neuroscience, 212, 159–179. https://doi.org/10.1016/j.neuroscience.2012.03.028

Lopez, Christophe, Lenggenhager, B., & Blanke, O. (2010). How vestibular stimulation interacts with illusory hand ownership. Consciousness and Cognition, 19(1), 33–47. https://doi.org/10.1016/j.concog.2009.12.003

Macauda, G., Bertolini, G., Palla, A., Straumann, D., Brugger, P., & Lenggenhager, B. (2015). Binding body and self in visuo-vestibular conflicts. The European Journal of Neuroscience, 41(6), 810–817. https://doi.org/10.1111/ejn.12809

Maister, L., Slater, M., Sanchez-Vives, M. V., & Tsakiris, M. (2015). Changing bodies changes minds: owning another body affects social cognition. Trends in Cognitive Sciences, 19(1), 6–12. https://doi.org/10.1016/j.tics.2014.11.001

McGeoch, P. D., Brang, D., Song, T., Lee, R. R., Huang, M., & Ramachandran, V. S. (2011). Xenomelia: a new right parietal lobe syndrome. Journal of Neurology, Neurosurgery, and Psychiatry, 82(12), 1314–1319. https://doi.org/10.1136/jnnp-2011-300224

McGonigle, D. J., Hänninen, R., Salenius, S., Hari, R., Frackowiak, R. S. J., & Frith, C. D. (2002). Whose arm is it anyway? An fMRI case study of supernumerary phantom limb. Brain: A Journal of Neurology, 125(Pt 6), 1265–1274.

Moseley, G. L. (2007). Using visual illusion to reduce at-level neuropathic pain in paraplegia. Pain, 130(3), 294–298. https://doi.org/10.1016/j.pain.2007.01.007

Moseley, G. L., Olthof, N., Venema, A., Don, S., Wijers, M., Gallace, A., & Spence, C. (2008). Psychologically induced cooling of a specific body part caused by the illusory ownership of an artificial counterpart. Proceedings of the National Academy of Sciences of the United States of America, 105(35), 13169–13173. https://doi.org/10.1073/pnas.0803768105

Moseley, P., Fernyhough, C., & Ellison, A. (2013). Auditory verbal hallucinations as atypical inner speech monitoring, and the potential of neurostimulation as a treatment option. Neuroscience and Biobehavioral Reviews, 37(10 Pt 2), 2794–2805. https://doi.org/10.1016/j.neubiorev.2013.10.001

Newport, R., & Preston, C. (2011). Disownership and disembodiment of the real limb without visuoproprioceptive mismatch. Cognitive Neuroscience, 2(3–4), 179–185. https://doi.org/10.1080/17588928.2011.565120

Nielsen, T. (1963). Volition: A New Experimental Approach. The Scandinavian Journal of Psychology, 4, 6.

Ocklenburg, S., Rüther, N., Peterburs, J., Pinnow, M., & Güntürkün, O. (2011). Laterality in the rubber hand illusion. Laterality, 16(2), 174–187. https://doi.org/10.1080/13576500903483515

Oddo-Sommerfeld, S., Hänggi, J., Coletta, L., Skoruppa, S., Thiel, A., & Stirn, A. V. (2018). Brain activity elicited by viewing pictures of the own virtually amputated body predicts xenomelia. Neuropsychologia, 108, 135–146. https://doi.org/10.1016/j.neuropsychologia.2017.11.025

Ono, K., Mikami, Y., Fukuyama, H., & Mima, T. (2016). Motion-induced disturbance of auditory-motor synchronization and its modulation by transcranial direct current stimulation. The European Journal of Neuroscience, 43(4), 509–515. https://doi.org/10.1111/ejn.13135

Park, H.-D., Bernasconi, F., Bello-Ruiz, J., Pfeiffer, C., Salomon, R., & Blanke, O. (2016). Transient Modulations of Neural Responses to Heartbeats Covary with Bodily Self-Consciousness. The Journal of Neuroscience: The Official Journal of the Society for Neuroscience, 36(32), 8453–8460. https://doi.org/10.1523/JNEUROSCI.0311-16.2016

Pazzaglia, M., Haggard, P., Scivoletto, G., Molinari, M., & Lenggenhager, B. (2016). Pain and somatic sensation are transiently normalized by illusory body ownership in a patient with spinal cord injury. Restorative Neurology and Neuroscience, 34(4), 603–613. https://doi.org/10.3233/RNN-150611

Petkova, V. I., & Ehrsson, H. H. (2008). If I were you: perceptual illusion of body swapping. PLoS One, 3, e3832. https://doi.org/10.1371/journal.pone.0003832

Preston, C. (2013). The role of distance from the body and distance from the real hand in ownership and disownership during the rubber hand illusion. Acta Psychologica, 142(2), 177–183. https://doi.org/10.1016/j.actpsy.2012.12.005

R Development Core Team. (2017). R: A Language and Environment for Statistical Computing. Vienna, Austria; R Foundation for Statistical Computing. Retrieved from http://www.R-project.org

Ramachandran, V. S., & McGeoch, P. (2007). Can vestibular caloric stimulation be used to treat apotemnophilia? Medical Hypotheses, 69(2), 250–252. https://doi.org/10.1016/j.mehy.2006.12.013

Ratcliffe, N., & Newport, R. (2017). The Effect of Visual, Spatial and Temporal Manipulations on Embodiment and Action. Frontiers in Human Neuroscience, 11, 227. https://doi.org/10.3389/fnhum.2017.00227

Revelle, W. (2017). psych: Procedures for Psychological, Psychometric, and Personality Research (Version R package version 1.7.8). Evanston, Illinois: Northwestern University. Retrieved from https://CRAN.R-project.org/package=psych

Rode, G., Charles, N., Perenin, M. T., Vighetto, A., Trillet, M., & Aimard, G. (1992). Partial remission of hemiplegia and somatoparaphrenia through vestibular stimulation in a case of unilateral neglect. Cortex; a Journal Devoted to the Study of the Nervous System and Behavior, 28(2), 203–208.

Romano, D., Pfeiffer, C., Maravita, A., & Blanke, O. (2014). Illusory self-identification with an avatar reduces arousal responses to painful stimuli. Behavioural Brain Research, 261, 275–281. https://doi.org/10.1016/j.bbr.2013.12.049

Salomon, R., Lim, M., Pfeiffer, C., Gassert, R., & Blanke, O. (2013). Full body illusion is associated with widespread skin temperature reduction. Frontiers in Behavioral Neuroscience, 7, 65. https://doi.org/10.3389/fnbeh.2013.00065

Serino, A., Alsmith, A., Costantini, M., Mandrigin, A., Tajadura-Jimenez, A., & Lopez, C. (2013). Bodily ownership and self-location: Components of bodily self-consciousness. Consciousness and Cognition, 22(4), 1239–1252. https://doi.org/10.1016/j.concog.2013.08.013

Sierra, M., Baker, D., Medford, N., & David, A. S. (2005). Unpacking the depersonalization syndrome: an exploratory factor analysis on the Cambridge Depersonalization Scale. Psychological Medicine, 35(10), 1523–1532. https://doi.org/10.1017/S0033291705005325

Sierra, M., & David, A. S. (2011). Depersonalization: A selective impairment of self-awareness. Consciousness and Cognition, 20(1), 99–108. https://doi.org/10.1016/j.concog.2010.10.018

Slotema, C. W., Blom, J. D., van Lutterveld, R., Hoek, H. W., & Sommer, I. E. C. (2014). Review of the efficacy of transcranial magnetic stimulation for auditory verbal hallucinations. Biological Psychiatry, 76(2), 101–110. https://doi.org/10.1016/j.biopsych.2013.09.038

Smit, M., Kooistra, D. I., van der Ham, I. J. M., & Dijkerman, H. C. (2017). Laterality and body ownership: Effect of handedness on experience of the rubber hand illusion. Laterality: Asymmetries of Body, Brain and Cognition, 22(6), 703–724. https://doi.org/10.1080/1357650X.2016.1273940

Tsakiris, M., Costantini, M., & Haggard, P. (2008). The role of the right temporo-parietal junction in maintaining a coherent sense of one’s body. Neuropsychologia, 46(12), 3014–3018. https://doi.org/10.1016/j.neuropsychologia.2008.06.004

Vallar, G., & Ronchi, R. (2009). Somatoparaphrenia: a body delusion. A review of the neuropsychological literature. Experimental Brain Research, 192(3), 533–551. https://doi.org/10.1007/s00221-008-1562-y

